# Programmed cell senescence in the mouse developing spinal cord and notochord

**DOI:** 10.1101/2020.07.24.220368

**Authors:** Jorge Antolio Domínguez-Bautista, Pilar Sarah Acevo-Rodríguez, Susana Castro-Obregón

## Abstract

Programmed cell senescence is a cellular process that seems to contribute to morphogenesis during embryo development, in addition to cell proliferation, migration, differentiation and programmed cell death, and has been observed in evolutionary distant organisms like mammals, amphibians and fish. Programmed cell senescence is a phenotype similar to stress-induced cellular senescence, characterized by the expression of cell cycle inhibitors such as CDKN1A/p21, increased activity of a lysosomal enzyme with beta-galactosidase activity (coined senescence-associated beta-galactosidase) and, most importantly, secretion of growth factors, interleukins, chemokines, metalloproteases, etc., collectively known as a senescent-associated secretory phenotype that instructs surrounding tissue. How wide is the distribution of programmed cell senescence during mouse development and its specific mechanisms to shape the embryo are still poorly understood. Here, we investigated whether markers of programmed cell senescence are found in the developing mouse spinal cord and notochord. We found discrete areas and developmental windows with high senescence-associated beta galactosidase in both spinal cord and notochord; expression of CDKN1A/p21 was documented in epithelial cells of the spinal cord and the notochord. Treatment of mice embryos developed ex-utero in the presence of the senolytic ABT-263 resulted in decrease senescence-associated beta-galactosidase activity and number of motoneurons. Our data suggest that several cell types undergo programmed cell senescence in developing spinal cord and notochord contributing to morphogenesis.

**Contribution to the Field Statement:** Cellular senescence is a state in which cells no longer divide but have the remarkable ability to secrete signaling molecules that alter the tissue where they reside. In adults, this state is typically induced by stress situations that cause DNA damage so cells with altered genome do not multiply. Senescent cells also form when a tissue is injured; they help to regenerate damaged tissue and contribute to wound healing. Phagocytic cells eliminate them when their function is done, having a transient existence. During vertebrate development some cells acquire a very similar phenotype, coined programmed cell senescence, and interestingly they have been found in regions that organize the pattern of development of some organs. How wide is the distribution of programmed cell senescence during development and how they help to shape the embryo are still poorly understood. We discovered in mice embryos different types of cells with senescent features located in particular regions of the developing nervous system: where motoneurons form and in a region that secrete molecules that instruct the embryo where different types of neurons will be created. We propose that programed cell senescence contributes to the morphogenesis of the nervous system.

## Introduction

Embryonic development is achieved by strongly coordinated mechanisms of cell migration, proliferation, differentiation and programmed cell death. Recently, programmed cell senescence was identified as an additional process that controls mouse development (Munoz-Espin et al., 2013; Storer et al., 2013). The presence of programmed cell senescence has also been reported during specific structures and windows of amphibian development, suggesting that programmed cell senescence contributes to vertebrate organogenesis and may have arisen in evolution as a developmental mechanism (Davaapil et al., 2017).

The development of the spinal cord and the differentiation of motoneurons are intensively studied due to its importance in organismal physiology and as in search of therapeutic targets for diseases such as amyotrophic lateral sclerosis. During the development of the spinal cord, the roof plate and the floor plate, located in the dorsal and ventral regions of the neural tube, regulate the dorsoventral patterning of different neuron populations by expressing WNT and BMP proteins dorsally and SHH ventrally (Chizhikov and Millen, 2004; Placzek and Briscoe, 2005). Interestingly, discrete populations of floor plate cells have been identified along the anteroposterior axis suggesting a variable mode of floor plate induction (Placzek and Briscoe, 2005). Adjacent to the ventral neural tube is located the notochord, a rod-like mesodermal structure that runs the anterior-posterior length, which becomes the rostrocaudal axis. The notochord is a source of developmental signals, such as the Hedgehog proteins, that play key roles in the patterning and proliferation of several organs (Corallo et al., 2015). As a result of signaling gradient of molecules secreted from the notochord and the floor plate, different progenitor cells located in the dorso-ventral axis of the neural tube give rise to V0-3 interneurons and motoneurons (Davis-Dusenbery et al., 2014).

In the developing spinal cord, blood vessels sprout from the perineural vascular plexus and invade the spinal cord at the ventral side. The motoneurons play an active role during blood vessel formation in the spinal cord by expressing vascular endothelial growth factor (VEGF) to allow blood vessel growth, but at the same time express a soluble VEGF receptor to titrate the availability of the growth factor in order to pattern the vasculature and block premature ingression of vessels by an attraction-repulsion mechanism (Himmels et al., 2017).

Cellular senescence is characterized by a durable exit from the cell cycle and acquisition of a flattened morphology, expansion of mitochondrial and lysosomal networks, a senescence-associated secretory phenotype (chemokines, cytokines, growth factors, metalloproteases, etc.), expression of cell cycle inhibitors as CDKN2A/p16, CDKN1A/p21 and/or p53, and a high level of senescence-associated-β-galactosidase activity (SA-β-gal). Senescent cells are metabolically active and influence the tissue microenvironment through their secretory phenotype (Czarkwiani and Yun, 2018). During development senescence contributes to tissue remodeling in structures such as the mesonephros, endolymphatic sac and apical ectodermal ridge in mice. Mechanistically, programmed cell senescence depends on the CDKN1A/p21 cell cycle inhibitor and is regulated by the TGF-β/Smad, PI3K/FOXO and ERK1/2 pathways (Munoz-Espin et al., 2013; Storer et al., 2013). In amphibians also TGF-β triggers programmed cell senescence, although is independent of ERK1/2 pathways (Davaapil et al., 2017). In contrast to other non-proliferating states as senescence or terminal differentiation, quiescence is a reversible cell cycle exit that takes place in cells that require a strict proliferation regime, such as stem cells (Sang et al., 2008; Sueda et al., 2019). Molecularly, the transcriptional repressor HES1 safeguards against irreversible cell cycle exit during cellular quiescence, and prevents senescence (Sang et al., 2008; Sueda et al., 2019).

Here we investigated whether there are cells undergoing programmed senescence during mammal spinal cord and notochord development. Our data show that in the roof plate, floor plate, motoneuron zone, and notochord there are indeed cells with SA-β-gal activity; nevertheless, only the notochord and endothelial cells in the spinal cord expressed also the cell cycle inhibitor CDKN1A/p21. To distinguish senescent cells from cell cycle arrested cells, we analyzed the expression of the quiescence marker HES1. Interestingly, cells at the roof and floor plates also express HES1. Additionally, treatment of mouse embryos with the senolytic drug ABT-263 resulted in decreased numbers of motoneurons. It is tempting to speculate that programmed cell senescence cells cooperate with programmed cell death to regulate motoneuronal population during spinal cord development.

## Materials and methods

### Timed mating and open-book preparations of spinal cords

Mice used in the present study were handled and cared according to the animal care and ethics legislation. All procedures were approved by the Internal Committee of Care and Use of Laboratory Animals of the Institute of Cell Physiology-UNAM (IFC-SCO51-18). Mice had *ad libitum* access to water and food. Estrous cycle of female CD1 mice was monitored by vaginal lavage examination. Females in proestrus or estrus were placed in cages with fertile males overnight. The following morning (9:00 a.m.) was considered as embryonic day 0.5 (E0.5) if a vaginal plug was found. On days E10.5 to E14.5, dams were killed by cervical dislocation, and the uterus was washed in PBS. Embryos were dissected under a stereoscopic microscope. For open-book preparations, spinal cords were dissected from E12.5 to E14.5 embryos, opened dorsally and freed from meninges.

### SA-β-gal assay

SA-β-gal assay was performed as described elsewhere (Debacq-Chainiaux et al., 2009). Whole embryos from stages E10.5 and E11.5, or open book preparations from spinal cords from E12.5 to E14.5 were washed twice in PBS and fixed 5 min with 2% formaldehyde plus 0.2% glutaraldehyde in PBS at room temperature. Reaction was performed by incubating embryos at 37 °C overnight in reaction solution containing 1 mg/ml X-gal, 40 mM citric acid/sodium phosphate buffer pH 6.0, 5 mM potassium ferrocyanide, 5 mM potassium ferricyanide, 150 mM sodium chloride, and 2 mM Magnesium chloride. After SA-β-gal assay, spinal cords were fixed with 4% paraformaldehyde for 20 min. Some samples were used to perform immunofluorescence after fixation with 4% paraformaldehyde.

### Immunofluorescence

Whole embryos or dissected spinal cords were fixed in PBS containing 4% paraformaldehyde for 30 min at room temperature, washed twice with PBS, and cryoprotected overnight with 30% sucrose in PBS. Embryos were cryosectioned by embedding in Tissue-Tek® O.C.T., histological sections of 40 µm were obtained using a Leica Cryostat and mounted on Superfrost plus slides. Sections on slides were washed three times with PBS for 5 min each time followed by permeabilization with 0.5% Triton™ X-100 in PBS for 30 min at room temperature. Afterwards, tissue sections were blocked with 2% Bovine Serum Albumin in PBS for 1 h and then incubated overnight at 4 °C with the following antibodies: ISLET-1 (1:200, abcam, ab109517), HES1 (1:200, Santa Cruz, sc-25392), F4/80 (1:200, abcam, ab6640), BCL-X_L_ (1:50, Santa Cruz, sc-1690) and CDKN1A/p21 (1:100, abcam, ab109199). Next day, secondary antibodies Alexa Fluor® 488 anti-rat (Invitrogen, A21208), Alexa Fluor® 488 anti-mouse (Invitrogen, A11029), Alexa Fluor® 488 anti-rabbit (Invitrogen, A11034), Alexa Fluor® 594 anti-rabbit (Invitrogen, A11037), or Alexa Fluor® 594 anti-mouse (Invitrogen, A11032) were incubated at a dilution of 1:500 in 2% BSA for 1 h at room temperature. Slides were mounted with Fluoromount-G™, (Electron Microscopy Sciences, 1798425) and images were collected in a LSM800 (Zeiss) confocal microscope. All images were collected with 1 Airy Unit aperture of pinhole. The acquisition of SA-β-gal images by confocal microscopy was performed as described elsewhere (Levitsky et al., 2013).

### DNA-damage induced senescence in mouse embryonic fibroblasts and live dead assay

Mouse embryonic fibroblasts (MEFs) from passages between 3 and 5 were plated at a density of 6750 cells/cm^2^ in DMEM supplemented with 10% fetal bovine serum plus antibiotic/antimycotic (Life Technologies, 15240062). Next day, aqueous solution of etoposide was added to a final concentration of 120 μM for 2 h. Then, the medium was replaced with etoposide-free DMEM plus fetal bovine serum. Cells were cultured for 5 d at 37 °C and 5% CO_2_ without changing the medium. Then, cells were treated for 20 h with 10 µM ABT-263 or equivalent concentrations of DMSO (0.2%) as vehicle-control. After ABT-263 treatment cell viability was quantified using the LIVE/DEAD™ Viability/Cytotoxicity Kit (ThermoFischer, L3224). Briefly medium was replaced with serum-free medium containing 1 µM calcein and 1 µM ethidium homodimer for 25 min. Then, epifluorescent images were collected in live cells using 50% glycerol in PBS as mounting medium. Live or dead cells were quantified manually using the multi-point tool from Fiji software.

### Embryo culture

Mouse embryos of E12 stage were cultured in a roller bottle system using reported protocols (Takahashi and Osumi, 2010; Kalaskar and Lauderdale, 2014). Uterus from pregnant mice were dissected, washed twice in sterile PBS at 37 °C, and transferred to DMEM/F12 under a sterility hood. Embryos surrounded by the intact yolk sac were detached from the placenta and then freed from the yolk sac without separating it from the embryo. Dissected embryos were transferred to culture medium at 37 °C consisting of KnockOut DMEM (Life Technologies, 10829018) containing 10% KnockOut Serum Replacement (Life Technologies, 10828028), N-2 supplement (Life Technologies, 17502048), 2% bovine serum albumin suitable for cell culture (RMBIO BSA-BAF-25G), and antibiotic/antimycotic (Life Technologies, 15240062). Embryos were pre-cultured in rolling bottles in an incubator at 37 °C with continuous flow of 95% O_2_/5% CO_2_. Then, good heart rate and blood circulation was verified under a stereoscopic microscope for culturing the embryos for additional 15 h in the presence of the autophagy inhibitor spautin-1 (SIGMA, SML0440) or the senolytic ABT-263 (Chemgood, C-1009). Equivalent concentrations of DMSO were included as control conditions to exclude vehicle effect.

### Western blot

As LC3-II is not detectable in lysates of embryonic spinal cord by Western blot, it was enriched by preventing its degradation with chloroquine, added to culture medium to a final concentration of 100 µM during the last hour of embryo culture. The spinal cords were dissected with meninges and total lysates were obtained using a buffer consisting of 68.5 mM Tris HCl at pH 6.8, 2% SDS, and protease inhibitors cOmplete™ULTRA tablets (Sigma 5892791001). Lysate was cleared using a pestle sonicator for 10 s at 30% cycle, and centrifuging at 10 000 g for 10 min at room temperature. Protein was quantified using the DC™-protein assay kit (Bio-Rad, 5000112) using bovine serum albumin as standard. Protein was subjected to electrophoresis, transferred to a PVDF-FL membrane and analyzed using the LC3 antibody (MBL, PD014 at 1:2000 dilution overnight at 4 °C) or anti-β-actin (Santa Cruz, sc-47778 at 1:10 000 for 1 h at room temperature). Secondary antibodies (IRDye® 800CW Goat anti-Rabbit (LI-COR, 92632211)and IRDye® 680RD Goat anti-Mouse IgG (LI-COR, 92568070) were diluted 1:10 000 and incubated for 1 h at room temperature. Bands were visualized in an Odyssey ® CLx Imager, and quantified with the Image Studio Lite Version 5.2 software.

### Statistical analysis

Data are plotted as mean ± standard error of the mean. One-way ANOVA followed by Bonferroni’s multiple comparisons test was performed using GraphPad Prism version 6.01 for Windows, GraphPad Software. * P ≤ 0.05, ** P ≤ 0.01, *** P ≤ 0.001

## Results

### SA-β-gal activity is observed in the embryonic motoneurons and phagocytic cells of the developing spinal cord

To identify markers of cellular senescence in the developing spinal cord, colorimetric detection of SA-β-gal and immunostaining for CDKN1A/p21 were performed in mice embryos. Cross-sections of E10.5 and E11.5 embryos or open-book preparations of spinal cords from E12.5 to E14.5 embryos showed strong SA-β-gal staining in the floor plate (stages E11.5 to E14.5) and the motoneuron zone (Figure 1A and B). Interestingly, the enzymatic activity observed in the floor plate and motoneuron zone gradually decreased from the rostral region towards the caudal region in spinal cords from E12.5 to E14.5 (Figure 1B). Note that in the medulla oblongata (m.o.) there is a cluster of cells positive for SA-β-gal (white arrowheads, Figure 1B). In fact, these cells also express the motoneuron marker ISLET-1 similar to motoneurons in the spinal cord (s.c.), as shown in Figure 1D. Additionally, cross-sections of E10.5 and E11.5 embryos showed that the notochord also displays SA-β-gal activity (Figure 1A), which is consistent with a previous report analyzing the chicken notochord (Lorda-Diez et al., 2015).

**Figure 1.**
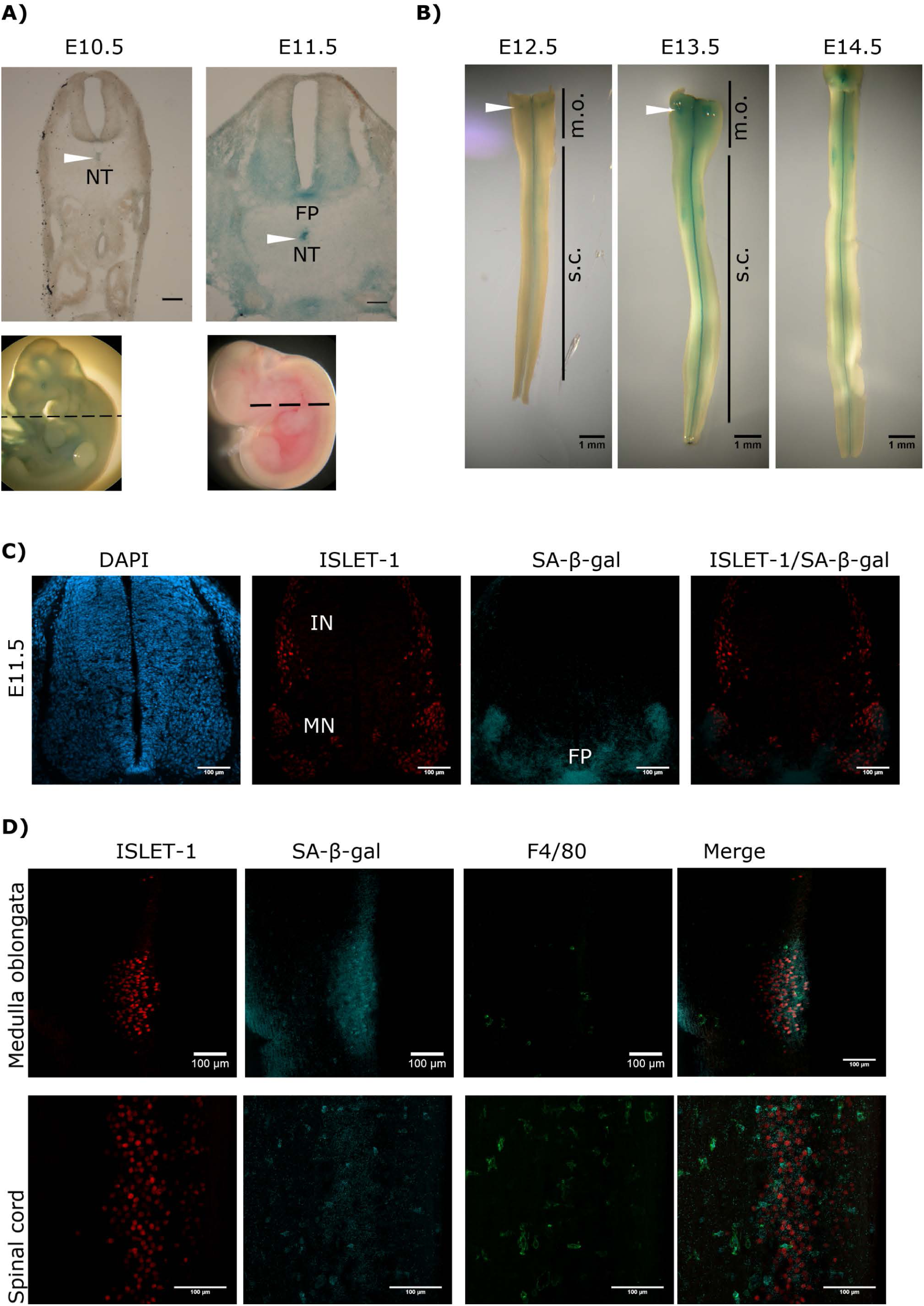
Transient and region-specific SA-β-gal activity in the mouse embryonic spinal cord and notochord. **(A)** Whole embryos from E10.5 and E11.5 were stained for SA-β-gal activity and cross-sectioned at the position indicated with dotted line in embryos shown in the lower row. Notice that the notochord (NT, white arrowheads), the floor plate (FP) and the ventral neural tube where motoneurons develop have high SA-β-gal activity at E11.5. Scale bars represent 100 µm. **(B)** Open-book preparations of spinal cords from E12.5 to E14.5 were subjected to SA-β-gal activity assay. SA-β-gal activity is strongly detected in the floor plate and at lower extent in the region corresponding to motoneurons of the spinal cord (s.c.). White arrowheads indicate SA-β-gal positive cells in the medulla oblongata (m.o.). **(C)** Whole E11.5 embryos were stained for SA-β-gal activity, thoracic sections were obtained and processed for immunodetection of ISLET-1. Confocal analysis shows that SA-β-gal activity overlaps with the motoneuron (MN) zone and the floor (FP) plate, while interneuron (IN) cells that also express ISLET-1, do not display SA-β-gal activity. Maximal projection is shown. Scale bars represent 100 µm. **(D)** Open-book spinal cord of stage E13.5 was stained for SA-β-gal activity and then subjected to whole-mount immunofluorescence against ISLET-1 and F4/80. Cells positive for ISLET-1 in the medulla oblongata (upper row) correspond to cells indicated with white arrowhead in Figure 1B. Detection of SA-β-gal activity by confocal microscopy combined with immunodetection of motoneurons (ISLET-1) and activated macrophages (F4/80) shows that these two cell types display SA-β-gal activity both in the medulla oblongata (upper row) and spinal cord (lower row). Maximal projections are shown. Scale bars represent 100 µm. Representative images are shown from at least 3 embryos or dissected spinal cords analyzed.

To demonstrate that the staining of SA-β-gal overlaps with the motoneuron zone, the motoneuron marker ISLET-1 was immunodetected in cross-sections of embryos previously stained for SA-β-gal and analyzed by confocal microscopy. As expected, the SA-β-gal activity detected by confocal microscopy overlaps with the motoneuron (MN) columns, but not with the interneurons (IN), which also express ISLET-1. Importantly, SA-β-gal activity was confirmed in the floor plate (FP, Figure 1C). Interestingly, we observed that indeed high SA-β-gal is observed in motoneurons located in the medulla oblongata (Figure 1D, upper row), and in the spinal cord (Figure 1D, lower row) in open-book preparations of E13.5 spinal cords stained for SA-β-gal and processed as whole mount for the immunodetection of ISLET-1. The possibility that SA-β-gal could be detected also in phagocytic cells was considered, given the occurrence of phagocytosis during development to clear apoptotic cells generated by programmed cell death, and possibly to clear also senescent cells. Since it has been described that macrophages have high lysosomal activity (Hall et al., 2017), we investigated whether some of the cells displaying SA-β-gal activity also express the activated macrophage/microglia marker F4/80. As shown in Figure 1D, some macrophages in the spinal cord indeed clearly showed strong SA-β-gal activity.

### The cell cycle inhibitor CDKN1A/p21 is not expressed in motoneurons but it is expressed in endothelial cells of the developing spinal cord and the notochord

To corroborate the senescent phenotype in the embryonic spinal cord and notochord, we analyzed whether the senescence marker CDKN1A/p21 was also expressed. First, we confirmed the specificity of our antibody by analyzing the apical ectodermal ridge, a structure known to express CDKN1A/p21 as part of the developmentally programmed cell senescence (Storer et al., 2013). Accordingly, cells in this structure indeed showed immunoreactivity against this antibody (Supplementary figure 1A). Then, cross-sections from E10.5 and E11.5 embryos were analyzed for the expression of CDKN1A/p21. Notably, CDKN1A/p21 was expressed in several parts of the embryos, including the spinal cord (Figure 2A) and the notochord, where it was located mainly in the cytoplasm (Figure 2B). We could not find motoneurons with a clear expression of CDKN1A/p21 in the spinal cord (Figure 2C), which was surprising since some of them had high SA-β-gal activity as shown before (Figures 1C and 1D). We identified as endothelial cells the subpopulation of cells in the spinal cord that expressed CDKN1A/p21 (Figure 2C). Endothelial cells in the medulla oblongata and in areas of the forelimb and thorax also expressed CDKN1A/p21 (Supplementary figure 1B and C). These results show that CDKN1A/p21 is expressed in endothelial cells of the spinal cord and other parts of the embryo during development. As cellular senescence is characterized by exit from the cell cycle and lack of cell division, we analyzed the expression of the proliferation marker Ki67. We found that only few cells along the central canal of the spinal cord express Ki67, none in the motoneuron zone (Figure 2D).

**Figure 2.**
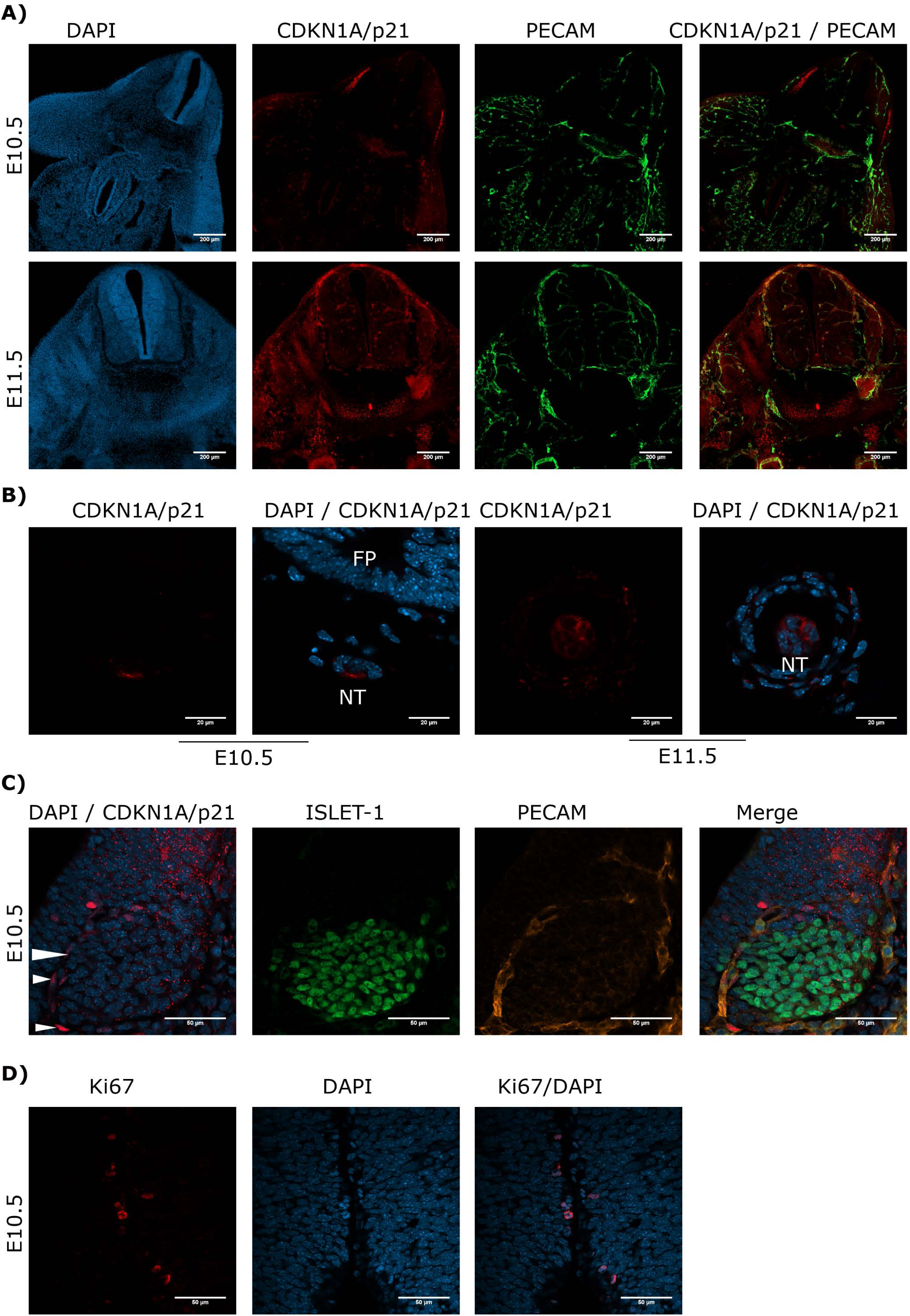
The cell cycle inhibitor CDKN1A/p21 is expressed in the mouse embryonic spinal cord and notochord. (**A**) Endothelial cells express CDKN1/p21. Brachial or cervical sections from E10.5 and E11.5 embryos, respectively, were processed for immunodetection of CDKN1A/p21 and PECAM. Embryos of the two stages show expression of CDKN1A/p21 in several organs, including the spinal cord. In the spinal cord, CDKN1A/p21 is expressed in endothelial cells labeled with PECAM. Single confocal planes are shown. Scale bars represent 200 µm. **(B)** CDKN1A/p21 is expressed in some notochord cells. Sections of embryos from stages E10.5 and E11.5 were processed as in A). Notice that CDKN1A/p21 is located in the cytoplasm of notochord (NT) cells, while it is absent in the floor plate (FP) from the E10.5 embryo. Single confocal planes are shown. Scale bars 20 µm. **(C)** CDKN1A/p21 is expressed in PECAM-positive endothelial cells localized in the vessels that surround the motoneuron column, but not in motoneurons themselves in the spinal cord of E10.5 embryos. White arrowheads indicate CDKN1A/p21-positive nuclei. Single confocal planes are shown. Scale bars 50 µm. **(D)** Only few cells along the central canal of the spinal cord of E10.5 embryos are proliferating, none in the motoneuron zone. Embryos from E10.5 stage were processed for immunofluorescence against Ki67 in cervical sections. Single confocal plane is shown. Scale bars represent 50 µm. Representative images are shown from at least 3 embryos analyzed.

### The floor and roof plates express the quiescence protein HES1

To distinguish quiescent cells from senescent cells, we analyzed the expression of the transcriptional repressor HES1, which prevents senescence establishment and irreversible cell cycle exit in quiescent cells (Sang et al., 2008; Sueda et al., 2019). Figure 3 shows that motoneurons identified by ISLET-1 expression did not expressed HES1. Intriguingly, several cells in the floor plate of spinal cords at E10.5 (Figure 3 upper panel) and E12.5 (and Figure 3 lower panel) expressed HES1, which is in accordance with a lack of expression of CDKN1A/p21 (Figure 2C, E10.5), but is in contrast with a strong activity of SA-β-gal in the floor plate (Figure 1A and B). The finding that the roof plate expressed HES1 at E12.5 is also in contrast with the report showing that the roof plate undergoes programmed cell senescence (Storer et al., 2013). Hence, in those areas there are senescent cells and quiescent cells.

**Figure 3.**
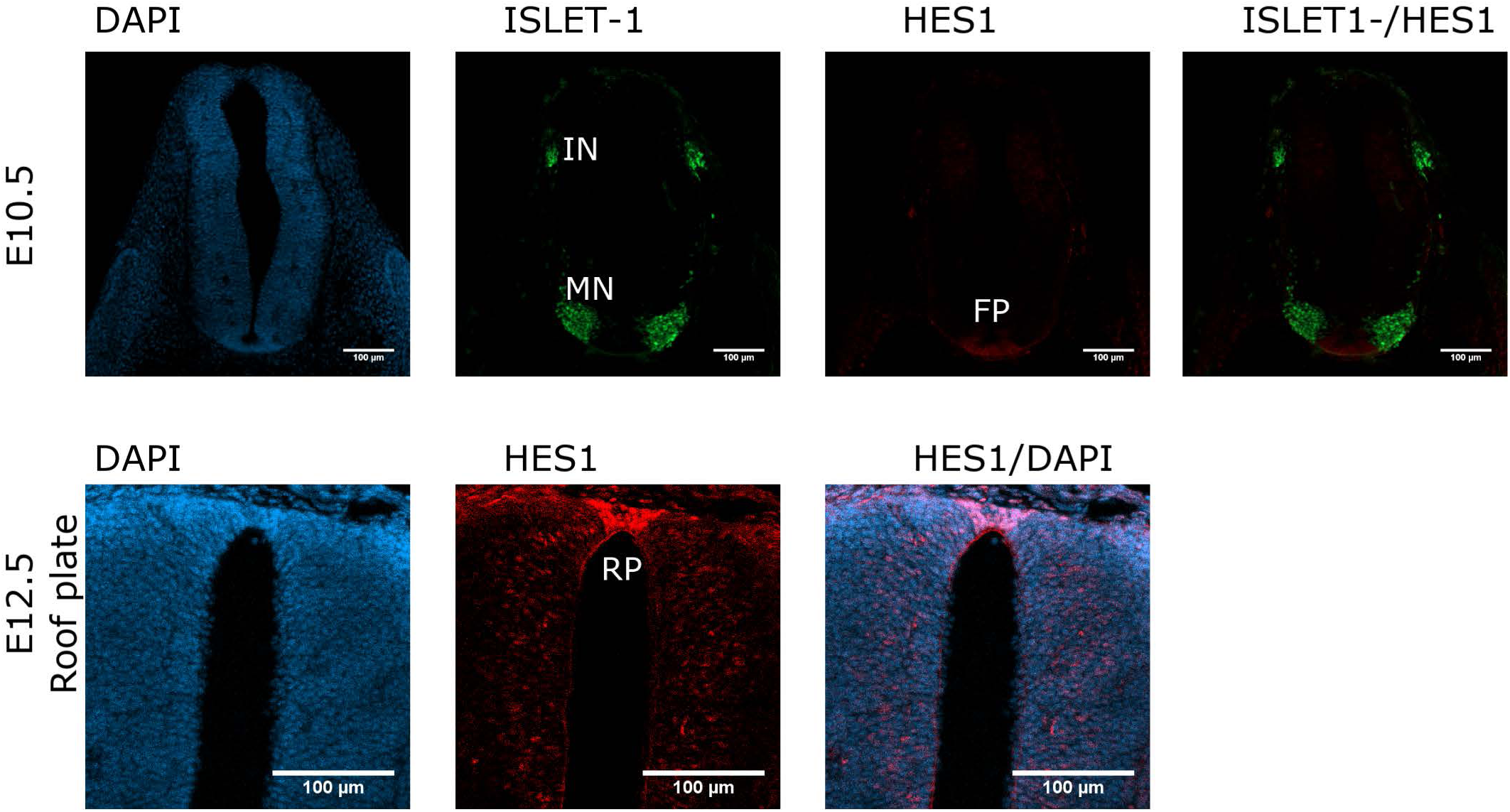
HES1 is expressed in some cells in the floor plate and roof plate in the developing spinal cord. Cross sections of E10.5 or E12.5 embryos (cervical or thoracic respectively) were analyzed for the expression of HES1 in the spinal cord. The floor plate of the E10.5 spinal cord contains cells with strong HES1 expression, but none was found in the motoneuron (MN) columns in E10.5 embryos. The roof plate of the E12.5 embryo also shows clear HES1 expression. Maximal projections of 4 confocal planes are shown. Scale bars 100 µm. Representative images are shown from at least 3 embryos analyzed.

### A subpopulation of embryonic motoneurons are sensitive to the senolytic drug ABT-263

Senescent cells are resistant to apoptosis thanks to an up regulation of members of the anti-apototic BCL-2 family, among other mechanisms. Accordingly, drugs that target BCL-X_L_ like proteins, such as ABT-263, have been developed as senolytics (Yosef et al., 2016). To determine whether the subpopulation of motoneurons with strong SA-β-gal activity we observed in the developing spinal cord were indeed senescent, we investigated whether they were sensitive to the senolytic ABT-263. To validate our tools, we compared the toxicity of ABT-263 in proliferative *vs*. senescent cells. By quantifying the number of live or dead cells, we confirmed that ABT-263 preferentially kills senescent fibroblasts (Supplementary figure 2). We then developed ex-utero mice embryos collected at stage E12 culturing them for 15 h in the presence of ABT-263 or vehicle. Mice embryos cultured with ABT-263 resulted in decreased activity of SA-β-gal in the spinal cord and apical ectodermal ridge (Supplementary figure 3A and B), confirming the elimination of programmed senescent cells. We verified the expression of BCL-X_L_ in the embryonic spinal cord (Figure 4A), including the motoneuron zone.

**Figure 4.**
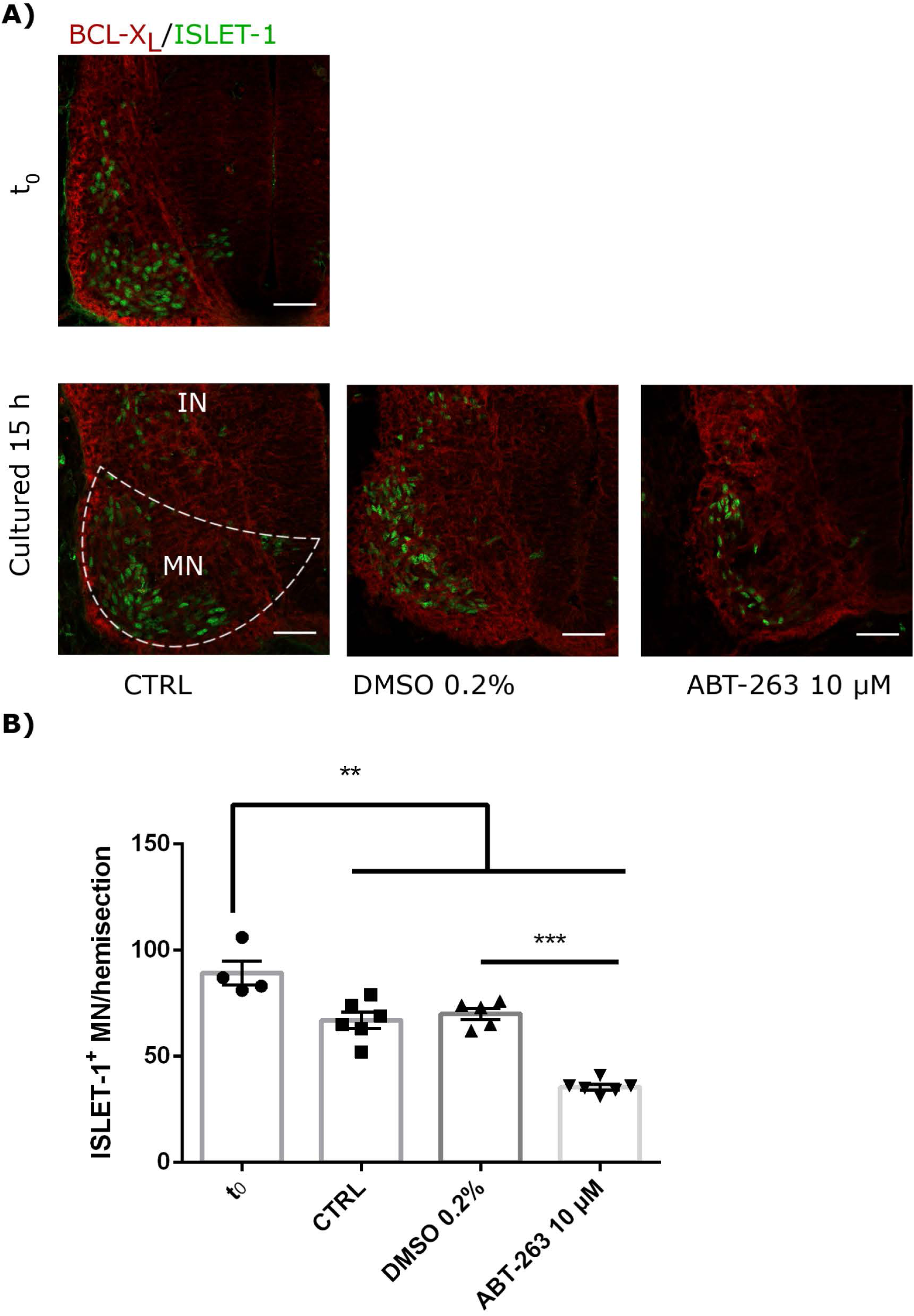
The senolytic ABT-263 promotes motoneurons cell death during development. **(A)** Cervical cross-sections of embryos at the beginning (t_0_) or after 15 h cultured with or without 10 µM ABT-263 were processed for immunofluorescence against BCL-X_L_ and ISLET1-1. Single confocal planes are shown. Scale bars 50 µm. **(B)** Quantification of motoneurons (ISLET-1) in embryos cultured with or without ABT-263. Each data point represents the number of cells quantified in the hemisection of a single embryo. At least 4 embryos were analyzed. Data are plotted as mean ± SEM significant differences were obtained from one-way ANOVA followed by Bonferroni’s multiple comparisons test. ** p ≤ 0.01; *** p ≤ 0.001

To analyze whether ABT-263 promotes motoneuron cell death, the number of ISLET-1 positive cells was quantified in embryos cultured with ABT-263 compared with control (CTRL) or vehicle (DMSO) conditions. As shown in Figure 4A and B, exposure to ABT-263 caused a statistically significant decrease in the number of motoneurons. To verify that the reduction in motoneuron number due to the ABT-263 treatment correlates with increased cell death, pyknotic nuclei were quantified in the motoneuron zone of the spinal cord and notochord. Indeed, treated embryos showed a higher amount of pyknotic nuclei in the motoneuron zone (Figure 5A upper row and B left). In comparison, the number of pyknotic nuclei in the notochord showed a trend to increase without reaching statistical significance (Figure 5A lower row and B right). These data show that during development, a subpopulation of motoneurons are sensitive to the senolytic ABT-263, thereby decreasing both the total number of motoneurons and SA-β-gal activity. We noticed that even though there was a broad expression of BCL-X_L_, only a subset of motoneurons was eliminated. These results suggest that a subpopulation of motoneurons undergo programmed cell senescence.

**Figure 5.**
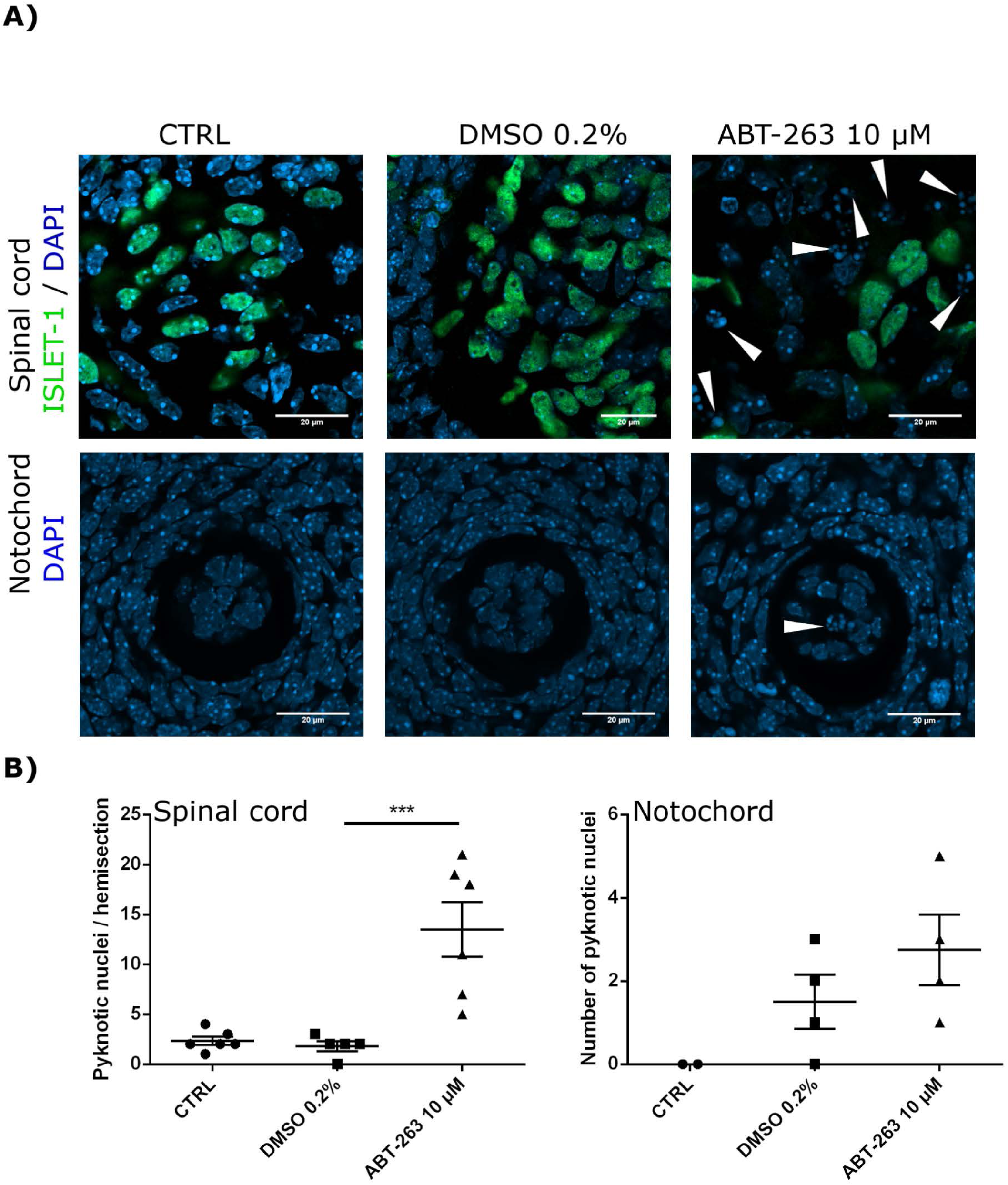
Embryos exposure to the senolytic ABT-263 increased pyknotic nuclei. **(A)** Confocal images of the spinal cord showing the motoneurons and nuclei (upper row), or the notochord (lower row) from embryos cultured with 10 µM ABT-263, vehicle (DMSO 0.2%) or control conditions. **(B)** Left: quantification of pyknotic nuclei in the spinal cord. Each data point represents the number of pyknotic nuclei in the motoneuron zone of a single embryo. At least 5 embryos were analyzed. Right: quantification of pyknotic nuclei in the notochord. Each data point represents the number of pyknotic nuclei in notochord. Number of embryos per group: CTRL n=2, DMSO n=4, ABT-263 n= 4. Data are plotted as mean ± SEM significant differences were obtained from one-way ANOVA followed by Bonferroni’s multiple comparisons test. *** p ≤ 0.001

During spinal cord development, roughly half of motoneurons are eliminated mainly, but not only, by apoptosis. We propose that a physiological function of motoneuronal programmed senescence could be to contribute to eliminate motoneurons during spinal cord development. Previous reports have shown that in addition to apoptosis other cellular processes can contribute to the programmed cell death of motoneurons, possibly cell death by autophagy (Sanchez-Carbente et al., 2005). To investigate whether autophagy signaling contributes to motoneurons cell death, mice embryos collected at stage E12 were cultured 15 h in the presence of the autophagy inhibitor spautin-1 (Liu et al., 2011). As occurs during normal development, the amount of ISLET-1-positive motoneurons decreased significantly during the course of the culture, reproducing *in vivo* programmed cell death. We found that autophagy inhibition with spautin-1 did not change the number of surviving motoneurons (Supplementary figure 4). These results show that pharmacological inhibition of autophagy during the developmental window of motoneurons programed cell death does not increase the amount of surviving ISLET-1 positive cells, suggesting that autophagic cell death does not contribute to establish the final number of motoneurons during spinal cord development.

In summary, our data show that embryonic motoneurons are insensitive to autophagy inhibition, but sensitive to the senolytic ABT-263, suggesting that programmed senescence contributes to the programmed elimination of motoneurons. It would be necessary to prevent motoneuron senescence establishment to test this idea.

## Discussion

Taking together, the data we present here suggest the presence of different cell types in the mouse embryonic spinal cord and notochord undergoing programmed cell senescence, as we found cells having strong SA-β-gal activity, CDKN1A/p21 expression and sensitivity to the senolytic ABT-263.

In the mouse notochord we found cells undergoing programmed cell senescence. In this structure cells had strong SA-β-gal activity (Figure 1A), which is consistent with a report studying chicken development (Lorda-Diez et al., 2015), and expressed CDKN1A/p21. Interestingly, CDKN1A/p21, was found in the cytosol in notochord cells (Figure 2B). While CDKN1A/p21 inhibits cell cycle when it is located in the nucleus, in the cytoplasm it has an anti-apoptotic function due to its ability to bind and inhibit the activity of proapoptotic proteins (Karimian et al., 2016). Additionally, the notochord from embryos cultured with ABT-263 showed reduced SA-β-gal activity and a trend to increase the amount of pyknotic nuclei compared with control or vehicle-treated embryos (Figure 5) suggesting that only a subpopulation of notochord cells become senescent during development. As the notochord acts as a signaling center for the differentiation of neuron populations of the spinal cord, a physiological role of senescent notochord cells is probably the contribution to the secretion of morphogens.

We propose that programmed cell senescence occurs in motoneurons during development of the spinal cord based on the following observations: there was a high SA-β-gal activity in the motoneuron zone during spinal cord development and we could identified SA-β-gal activity in ISLET-1 expressing cells (Figures 1C and 1D); SA-β-gal activity decreased in embryos developed ex-utero in the presence of the senolytic agent ABT-263 (Supplementary figure 3A), and the senolytic drug caused a significant decrease in the number of cervical motoneurons (Figure 4). This is an important finding because it has been previously reported that apart from apoptosis, other mechanisms may participate in the clearance of motoneurons during development (Sanchez-Carbente et al., 2005). We propose that a subpopulation of motoneurons become senescent either to secret morphogens or to be cleared by macrophages, contributing to the elimination of motoneurons during spinal cord development. In a similar way that senescent cells in the apical ectodermal ridge reintegrated in the tissue (Li et al., 2018), it could also occur that a subpopulation of senescent motoneurons continues to fully differentiate into postmitotic neurons and remain integrated in the tissue.

We did not find motoneurons expressing CDKN1A/p21, but it is still possible that they engage in programmed senescence in an event regulated by CDKN1B/p27 or CDKN2B/p15. When programmed senescence was identified as a mechanism that contributes to embryonic development, significant expression of CDKN1B/p27 or CDKN2B/p15 was detected, in addition to CDKN1A/p21, in the epithelia of mesonephric tubules and endolymphatic sac from E11.5 to E14.5; nevertheless, only the knockout of CDKN1A/p21 resulted in a partially altered SA-β-gal activity (Munoz-Espin et al., 2013). TGF-β is the signaling molecule that controls programmed senescence (Munoz-Espin et al., 2013), but it also promotes the activation of cell cycle inhibitors as a pre-requisite for neurogenesis (Casari et al., 2014). In fact, Smad3 overexpression in embryonic chick spinal cord resulted in upregulation of CDKN1B/p27, with consequent cell cycle exit and enhancement to neuronal differentiation (Garcia-Campmany and Marti, 2007). Considering that the cell cycle inhibitor CDKN1B/p27 has been linked to senescence in the context of cancer pathogenesis (Lin et al., 2010; Flores et al., 2014; Revandkar et al., 2016; Seo et al., 2018), it could also mediate motoneuronal programmed cell senescence.

To confirm a developmental role of senescent motoneurons, we should be able to prevent the acquisition of the senescent phenotype and observe a disruption of the developmental pattern of the neural tube, or an increase in the total number of motoneurons. Currently this experiment is not possible, since TGF-β is the only signaling pathway described to induce programmed senescence (Munoz-Espin et al., 2013) and it has also a crucial role in neural tube development. So, is we inhibit TGF-β pathway we won’t be able to distinguish the developmental role of TGF-β from the role of senescent cells *per se*.

Endothelial cells of the spinal cord perhaps undergo programmed cell senescence, as they expressed CDKN1A/p21 (Figures 2A and C); however, this result needs cautious interpretation. Unlike the programmed senescence that occurs in transient structures that are eliminated during development, such as the apical ectodermal ridge (Storer et al., 2013) or the mesonephros (Munoz-Espin et al., 2013), endothelial cells remain in the blood vessels where they become fundamental components. Consistent with our data, previous studies have shown that during endothelial cell differentiation two markers of senescence are observed: CDKN1A/p21, that mediates the cell cycle arrest necessary for endothelial cell differentiation (Zeng et al., 2006; Marcelo et al., 2013), and SA-β-gal activity, that was shown to occur during differentiation of human prostate epithelial cells (Untergasser et al., 2003). A report that elegantly linked the concepts of senescence and differentiation was published recently (Li et al., 2018). By using genetic lineage tracing, Li and colleagues showed that in the apical ectodermal ridge at mid-stage (E10.5-E13.5) there is a population of CDKN1A/p21-positive and SA-β-gal-positive cells (*i*.*e*., senescent cells). Interestingly, by late-stage (E15.5-E16.5) or after birth, a subset of the previously senescent cells re-entered the cell cycle, proliferated, exhibited epithelial fate and contributed to tissues (Li et al., 2018). Thus, endothelial cells that express CDKN1A/p21 in the developing spinal cord may engage in a senescent program and nevertheless form the blood vessels later in development.

Although senescent cells and quiescent cells seem to be present in same regions, SA-β-gal activity could occur in quiescent cells expressing HES1, as we observed that SA-β-gal activity in the floor plate (Figures 1A and B) and roof plate (see Supplementary figure 3A) overlap with the quiescence marker HES1 (Figure 3A), but not with CDKN1A/p21 (Figure 2B), consistent with previous work (Baek et al., 2006). To determine whether indeed the same cell expressing HES1 had high SA-β-gal, we attempted to detect both HES1 and SA-β-gal in the same preparations but this resulted in strong background and nonspecific binding of the HES1 antibody, probably caused by the fixation protocol for SA-β-gal. This observation is consistent with reports showing that SA-β-gal is detectable in non-senescent cells such as the yolk sac, visceral endoderm, and macrophages (Cristofalo, 2005; Yang and Hu, 2005; Huang and Rivera-Perez, 2014; Hall et al., 2017). The activation of SA-β-gal is also caused by cellular differentiation induced by TGF-β (Untergasser et al., 2003). Additionally, a recent report in the developing avian retina found that SA-β-gal activity correlated with CDKN1A/p21 expression, but authors concluded that SA-β-gal activity and CDKN1A/p21 expression occurred independently of programmed senescence because SA-β-gal activity did not correlate with the chronotopographical distribution of apoptotic cells (de Mera-Rodriguez et al., 2019). Nevertheless, apoptosis is not necessarily simultaneous with the senescent phenotype, nor the only fate of programmed senescent cells. Previous articles showed SA-β-gal in the neural tube, and concluded that programmed senescence occurs in the roof plate of the spinal cord (Munoz-Espin et al., 2013; Storer et al., 2013; Czarkwiani and Yun, 2018). Indeed, CDKN1A/p21-positive cells were detected in the closing neural tube (Storer et al., 2013), but expression of CDKN1A/p21 or other cell cycle inhibitor was not shown in the roof plate. Thus, we consider here that in addition to senescence, quiescence can be accompanied by SA-β-gal in the roof plate.

In summary, we propose that 1) SA-β-gal activation may occur during the quiescent phenotype of the floor and roof plates, 2) endothelial cells are probably engaged in a phenomenon of reversible senescence, 3) a subpopulation of notochord cells become senescent and secrete morphogens and 4) motoneurons programmed senescence could contribute to the elimination of motoneurons during spinal cord development. Further experiments will be needed to test these hypotheses.

## Author Contributions

JDB and SCO contributed to conception and designed of the study; JDB performed most of the experiments and wrote the first draft of the manuscript. PAR acquired data. All authors contributed to manuscripts revision, read and approved the submitted version of the manuscript and are accountable for the content of the work.

## Acknowledgements

We thank the following persons for their support during the course of this project: Dr. Beatriz Aguilar-Maldonado (technical assistance), Dr. Claudia Rivera-Cerecedo (Animal Facility), Dr. Ruth Rincón-Heredia and Dr. Abraham Rosas-Arellano (Microscopy facility), Eng. Aurey Galván Lobato and Eng. Manuel Ortínez Benavides (Equipment Maintenance Workshop). Data in this work is part of PAR doctoral dissertation in the Posgrado en Ciencias Bioquímicas de la Universidad Nacional Autónoma de México.

## Abbreviations

ANOVA: Analysis of Variance
BCL-X_L_: B-cell lymphoma-extra large
CDKN1A/p21: Cyclin-dependent kinase inhibitor 1A
CDKN1B/p27: Cyclin-dependent kinase inhibitor 1B
CDKN2A/p16: Cyclin-dependent kinase inhibitor 2A
CDKN2B/p15: Cyclin-dependent kinase inhibitor 2B
CQ: Chloroquine
CTRL: Control
DMEM: Dulbecco’s Modified Eagle Medium
DMEM/F12: Dulbecco’s Modified Eagle Medium/Nutrient Mixture F-12
DMSO: Dimethyl Sulfoxide
ERK1/2: Extracellular signal–Regulated Kinases 1 and 2
FP: Floor plate
IN: Interneurons
LC3: Microtubule-associated protein 1A/1B-light chain 3
MEFs: Mouse Embryonic Fibroblasts
MN: Motoneurons
PBS: Phosphate-Buffered Saline
PECAM: Platelet Endothelial Cell Adhesion Molecule
PI3K/FOXO: Phosphoinositide-3-Kinase/Forkhead box O
RP: Roof plate
SA-β-gal: Senescence-associated beta-galactosidase
SDS: Sodium Dodecyl Sulfate
SEM: Standard Error of the Mean
TGF-β: Transforming Growth Factor beta

## Supplementary Material

**Supplementary figure 1. The cell cycle inhibitor CDKN1A/p21 is expressed in endothelial cells of the medulla oblongata and forelimb. (A)** The specificity of the CDKN1A/p21 antibody was validated by analyzing the expression of CDKN1A/p21 in the apical ectodermal ridge of an E12.5 embryo. The section area is indicated on the diagram on the right. Single confocal planes are shown. Scale bars represent 20 µm. **(B)** Cross-section analysis of the medulla oblongata from an E12 embryo as indicated on the right. Confocal microscopy shows that CDKN1A/p21 is expressed in endothelial (PECAM-positive) cells. Upper panel low magnification, lower panel higher magnification. Single confocal planes are shown. Scale bars represent 100 µm or 20 µm. **(C)** Expression of CDKN1A/p21 in endothelial cells of the forelimb and thorax of an E13.5 embryo. Single confocal planes are shown. Scale bars 100 µm.

**Supplementary figure 2. *In vitro* validation of the senolytic effect of ABT-263**. Proliferative (non-senescent) and senescent MEFs were treated for 20 h with ABT-263 at the indicated concentrations and the cytotoxic effect was determined quantifying live and dead cells. Since 5 µM ABT-263 induced senescent fibroblasts death concomitant with a reduction of cells alive. We noticed a decrease in the number of live non-senescent MEFs with 10 µM ABT-263, although the number of dead cells was not affected. Each group was cultured in triplicate wells, and five random fields in each well were used to count the number of live or dead cells. Data are plotted as mean ± SEM significant differences were obtained from one-way ANOVA followed by Bonferroni’s multiple comparisons test. * p ≤ 0.05, ** p ≤ 0.01

**Supplementary figure 3. SA-β-gal activity in the apical ectodermal ridge and spinal cord was reduced after treatment with ABT-263. (A)** Embryos of the E12 stage were cultured in the presence of 5 µM ABT-263, vehicle (DMSO 0.2%) or control for 15 h and stained for SA-β-gal and cross-sectioned. The treatment with ABT-263 resulted in a decrease of SA-β-gal activity in the motoneuron zone and floor plate. **(B)** Embryos of the E12 stage were cultured as in A), and images of the apical ectodermal ridges were collected. Representative results from 3 embryos analyzed.

**Supplementary figure 4. Pharmacological inhibition of autophagy does not alter the amount of motoneurons. (A)** Treatment with spautin-1 inhibits autophagy, as it results in a decrease of LC3-II levels compared to control (CTRL) or vehicle (DMSO). Western blot to detect indicated proteins from total lysates obtained from embryonic spinal cords of E12 embryos cultured with spautin-1, vehicle or control for 15 h. LC3-II level was normalized detecting β-actin as loading control. As a reference to distinguish LC3-II, a lysate from mouse embryonic fibroblasts (MEFs) cultured with chloroquine (CQ) was also loaded. **(B)** ISLET-1 positive motoneurons were quantified in cervical cross-sections from embryos dissected at E12 and developed ex-utero for 15 h with indicated treatments. The results of four independent embryo cultures are shown, with at least 2 embryos per condition per culture. Each data point represents the number of cells quantified in the hemisection of a single embryo. The number of motoneurons significantly decreased in all conditions after culture, as occurs during this stage of embryo development, but pharmacologic inhibition of autophagy with spautin-1 did not affect the amount of surviving motoneurons. Data are plotted as mean ± SEM, significant differences were obtained from one-way ANOVA followed by Bonferroni’s multiple comparisons test. * p ≤ 0.05

