## Supplementary figures for "Programmed cell senescence in the mouse developing spinal cord and notochord"

**Supplementary figure 4. Pharmacological inhibition of autophagy does not alter the amount of motoneurons. (A)** Treatment with spautin-1 inhibits autophagy, as it results in a decrease of LC3-II levels compared to control (CTRL) or vehicle (DMSO). Western blot to detect indicated proteins from total lysates obtained from embryonic spinal cords of E12 embryos cultured with spautin-1, vehicle or control for 15 h. LC3-II level was normalized detecting β-actin as loading control. As a reference to distinguish LC3-II, a lysate from mouse embryonic fibroblasts (MEFs) cultured with chloroquine (CQ) was also loaded. **(B)** ISLET-1 positive motoneurons were quantified in cervical cross-sections from embryos dissected at E12 and developed ex-utero for 15 h with indicated treatments. The results of four independent embryo cultures are shown, with at least 2 embryos per condition per culture. Each data point represents the number of cells quantified in the hemisection of a single embryo. The number of motoneurons significantly decreased in all conditions after culture, as occurs during this stage of embryo development, but pharmacologic inhibition of autophagy with spautin-1 did not affect the amount of surviving motoneurons. Data are plotted as mean ± SEM, significant differences were obtained from one-way ANOVA followed by Bonferroni's multiple comparisons test. * $p \leq 0.05$

Supplementary figure 1

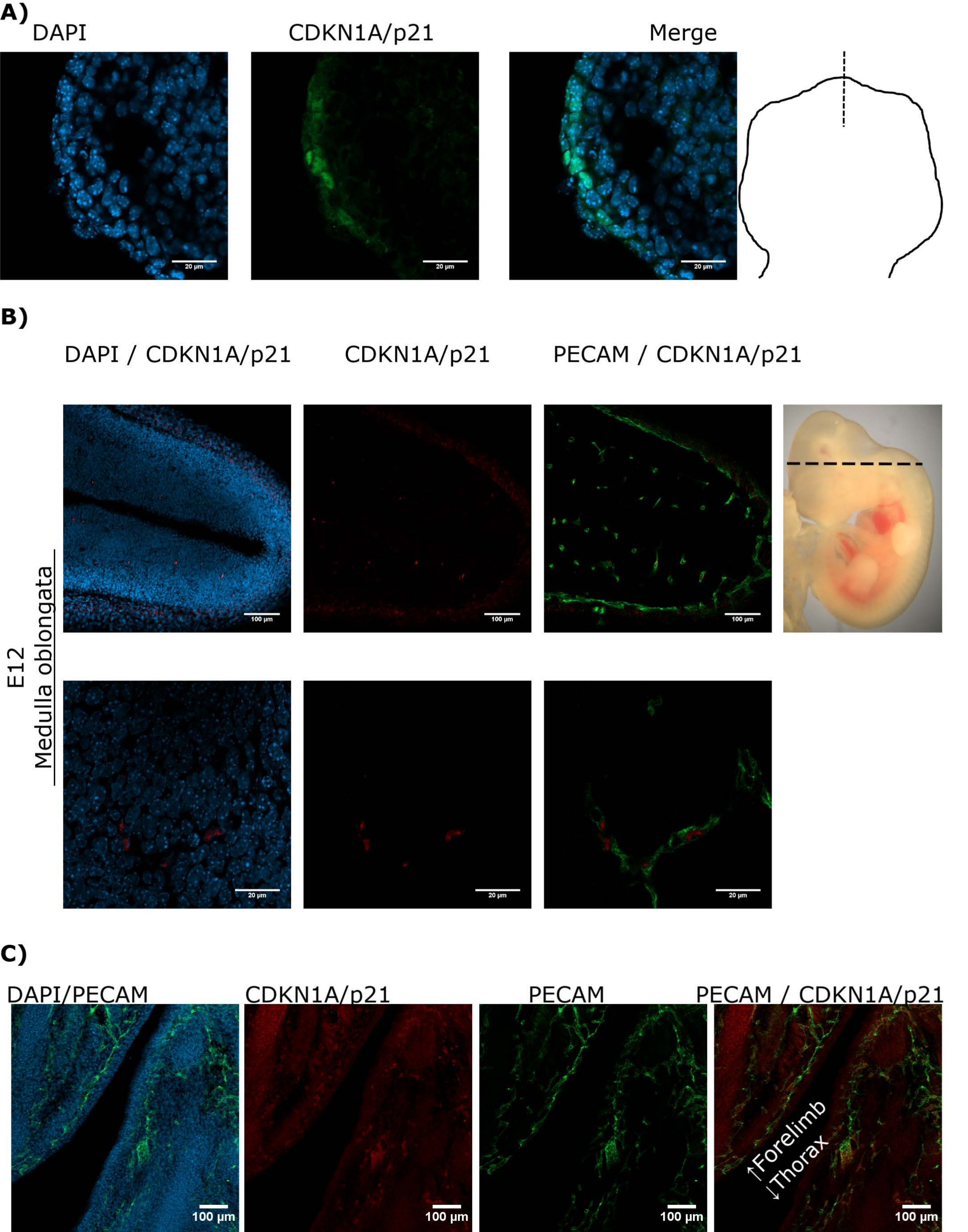

### Supplementary figure 2

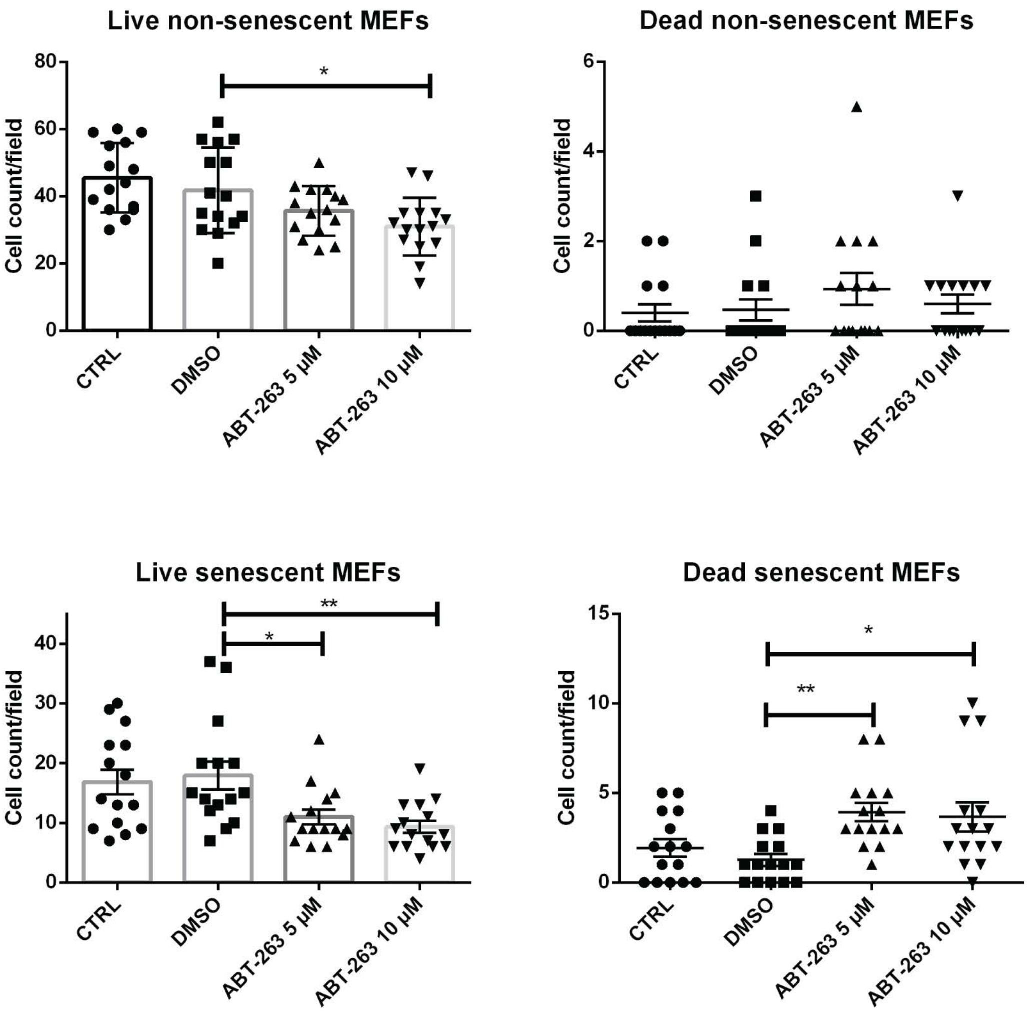

Supplementary figure 3

**A)**

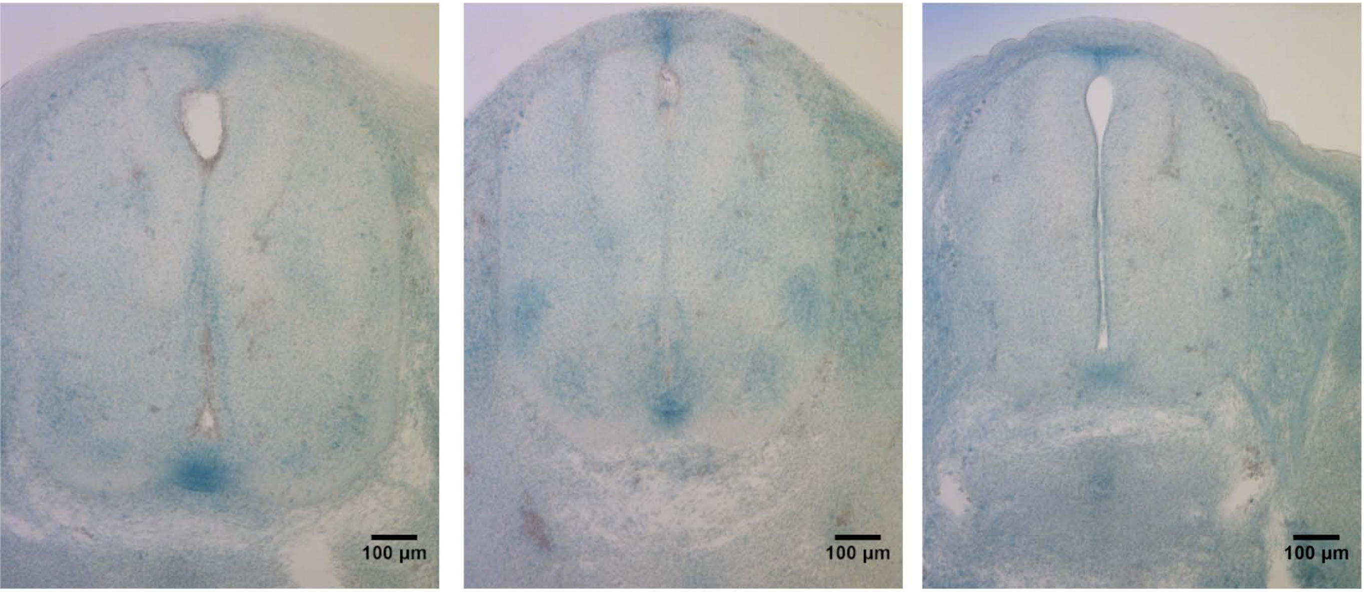


CTRL DMSO ABT-263 5 µM

**B)**

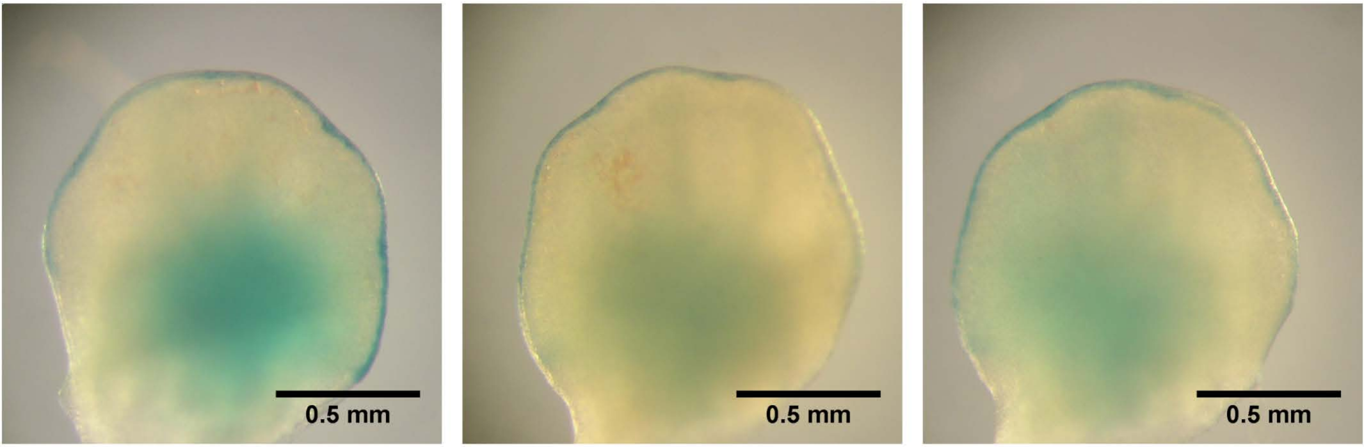


CTRL DMSO ABT-263 5 µM

Supplementary figure 4

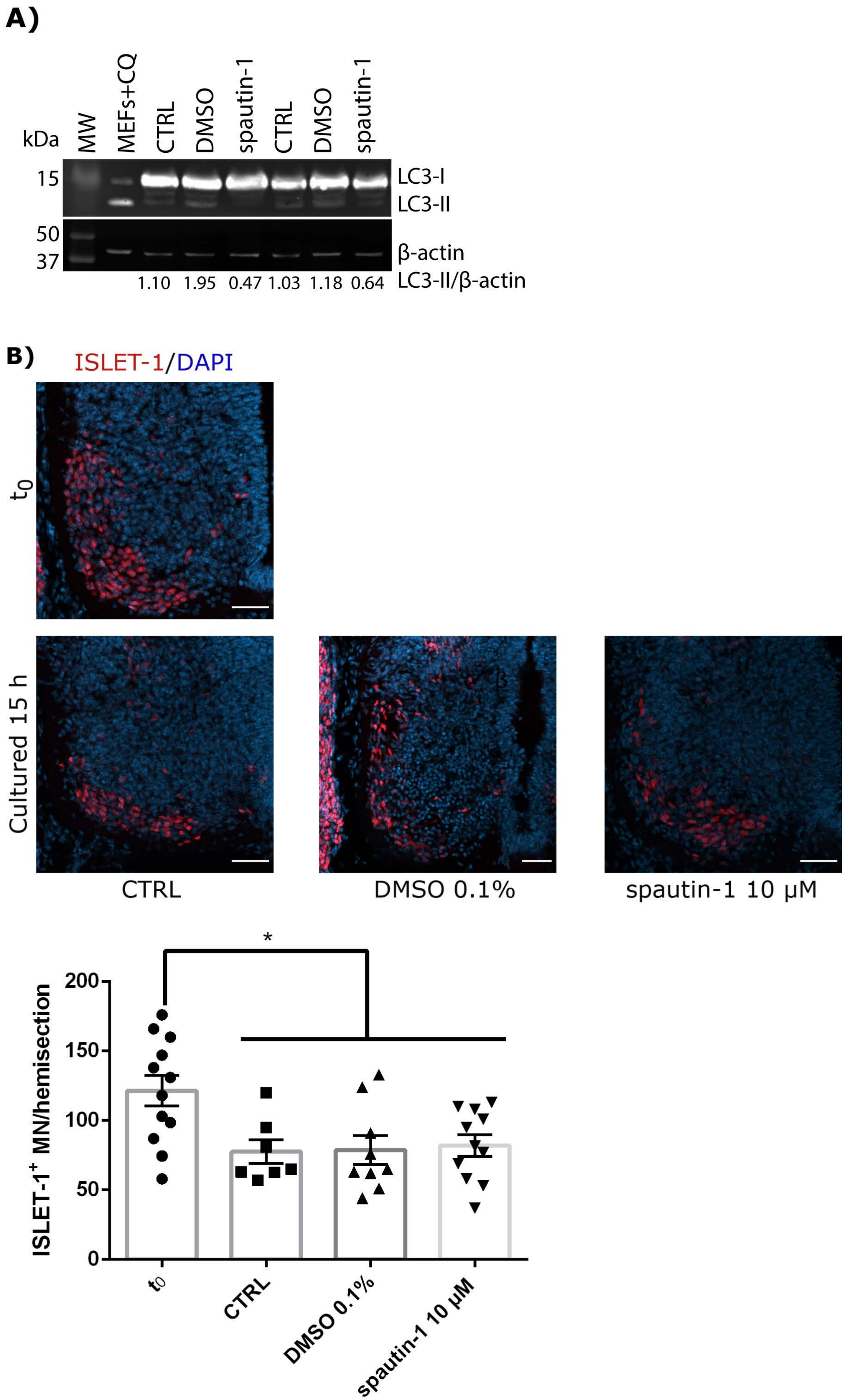
